# Hair cells in the cochlea must tune resonant modes to the edge of instability without destabilizing collective modes

**DOI:** 10.1101/2024.07.19.604330

**Authors:** Asheesh S. Momi, Michael C. Abbott, Julian Rubinfien, Benjamin B. Machta, Isabella R. Graf

## Abstract

Sound produces surface waves along the cochlea’s basilar membrane. To achieve the ear’s astonishing frequency resolution and sensitivity to faint sounds, dissipation in the cochlea must be canceled via active processes in hair cells, effectively bringing the cochlea to the edge of instability. But how can the cochlea be globally tuned to the edge of instability with only local feedback? To address this question, we use a discretized version of a standard model of basilar membrane dynamics, but with an explicit contribution from active processes in hair cells. Surprisingly, we find the basilar membrane supports two qualitatively distinct sets of modes: a continuum of *localized* modes and a small number of collective *extended* modes. Localized modes sharply peak at their resonant position and are largely uncoupled. As a result, they can be amplified almost independently from each other by local hair cells via feedback reminiscent of self-organized criticality. However, this amplification can destabilize the collective extended modes; avoiding such instabilities places limits on possible molecular mechanisms for active feedback in hair cells. Our work illuminates how and under what conditions individual hair cells can collectively create a critical cochlea.

## Introduction

The human cochlea is a spiral-shaped organ in the inner ear, where sound is converted into electrical signals. The cochlea can detect sounds with frequencies across three orders of magnitude (20–20 000Hz) and a trillion-fold range in power (0–130dB), down to air vibrations on the order of an angstrom. After entering the cochlea, sound waves become surface waves along the basilar membrane (BM), depositing most incident energy in a frequency-specific location [1].

Dissipation in the cochlea, through friction and viscous loss, limits frequency resolution and sensitivity. To counter dissipation, the cochlea contains active force-generating mechanisms [2–5]. Active processes are performed by hair cells, small sensory structures that line the BM. For overly strong hair cell activity, the BM becomes unstable to spontaneous oscillations. When activity almost cancels friction, the cochlea is highly sensitive to weak amplitude signals, and frequency selectivity is high. This barely-stable regime is thus ideal for sound processing, but appears to require fine-tuning. Here we ask how hair cells can find this operating regime.

Past models have provided insight into possible mechanisms for tuning individual hair cells [6–8]. In particular, that work studied how single hair cells can find a Hopf bifurcation, a transition between a stable and an unstable oscillatory regime. The discussion often focuses on bullfrog hearing where there is no cochlea, and hair cells act as relatively independent mechanical oscillators. Conversely, Models of the mammalian cochlea typically ignore tuning and operate under the assumption that active processes have globally reduced friction to near zero [9].

In this work, we argue that assuming that hair cells cancel friction for all frequencies and positions is neither necessary nor feasible, and we instead seek to understand how they can find an operating region with the desired properties of the dissipation-free state. Friction only dominates the dynamics of the cochlea precisely at resonance because passive mechanics are underdamped [10]. Furthermore, individual hair cells are small mechanical perturbations to the overall dynamics and their contribution to the non-dissipative mechanics is likely inconsequential. We thus focus on the role of hair cell activity in reducing friction in a manner which is local in both space and frequency. We show that, with some interesting caveats, this is sufficient to bring the cochlea to a line of Hopf bifurcations where every frequency is nearly critical at a specific location [11, 12].

Towards this end we expand on an established model for the dynamics of the cochlea, by explicitly adding mechanical contributions from active processes in hair cells. We assume that hair cells detect the displacement of the BM and respond by exerting forces via a fast linear response kernel. Each hair cell can also slowly adjust the strength of its active processes to find the global operating regime.

Perhaps surprisingly, this model of the cochlea contains two distinct types of modes. The first type, which we term *localized* modes, strongly peak at particular resonant positions. The second type instead have energy throughout the cochlea, reminiscent of standing waves, and we term them *extended* modes. While tuning activity is generally able to bring the localized modes to the edge of instability, the extended modes become unstable for many plausible forms of active processes. For suitable forms of active processes, we further propose a mechanism reminiscent of self-organized criticality [13, 14], which tunes the local activity strength to the edge of instability.

## Results

The passive part of our model of the cochlea has three components: the fluid which moves along the cochlea, the oval window by which the middle ear pushes this fluid, and the elastic basilar membrane which separates the cochlea into two compartments. We introduce each of these in turn and examine the resulting mode structure, before focusing on the active component, hair cells.

### Wave equation for the cochlea

In a widely used approximation [1], the cochlea consists of two fluid-filled compartments, the scala tympani and the scala vestibuli, which are separated by the BM (Fig. 1). Sound creates a fluid flux, coupled to a change in pressure described by force balance:

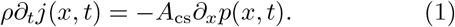

Here, *j*(*x, t*) is the difference in volume current between the lower and upper compartment, *p*(*x, t*) is the pressure difference, *A*_*cs*_ is the average cross-sectional area of a cochlear compartment, and *ρ* is the density of water. The fluid flux propagates down the cochlea, creating a displacement of the BM, *h*(*x, t*), which we call its height. The fluid flux obeys a continuity equation,

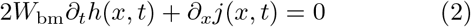

where *W*_*bm*_ is the width of the BM. We can eliminate *j*(*x, t*) from Eq. 1 and 2 to arrive at a modified wave equation relating height and pressure [1, 9, 15],

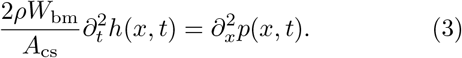

Sound enters the cochlea through the oval window, which connects to the middle ear and, in turn, the ear canal. We follow Ref. [15] for the boundary conditions at the base of the cochlea (*x* = 0), where the lateral displacement of the oval window *d*_ow_(*t*) creates a flux of fluid, and eventually an equal and opposite flux in the scala tympani’s round window. Via Eq. 1, this leads to a pressure gradient:

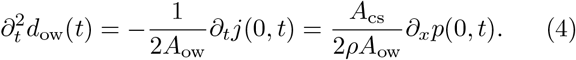

Here *A*_*ow*_ is the area of the oval window. The oval window itself acts as a damped harmonic oscillator:

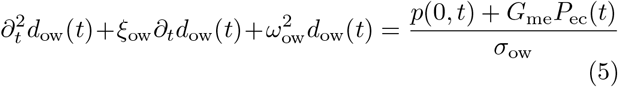

where *ξ*_ow_ is its dampening constant, *ω*_ow_ the middle ear resonance, *P*_ec_(*t*) the pressure in the ear canal, *G*_me_ the gain of the middle ear, and *σ*_ow_ the mass per area of the oval window. At the cochlea’s apex (*x* = L), a gap in the basilar membrane (the helicotrema) suggests zero pressure difference [15]:

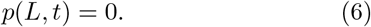

**Figure 1.**
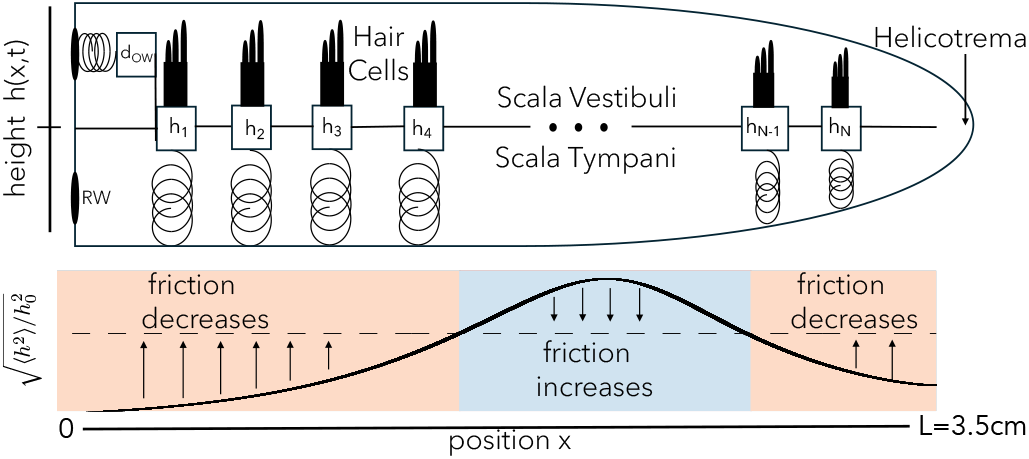
Schematic of the model. Above, we unroll the spiral-shaped cochlea into a model of two fluid-filled chambers partitioned by the basilar membrane (BM). Pressure waves in the fluid are accompanied by BM displacement *h*(*x, t*), which we model as a set of *N* damped harmonic oscillators, each with active driving from a hair cell. Sound input from the middle ear is via the displacement *d*_*ow*_(*t*) of the oval window (OW) and a corresponding flux at the cochlea’s base, which due to fluid incompressibility causes an equal and opposite flux at the round window (RW). The pressure difference at the helicotrema at *x* = *L* is set to zero. Below is a sketch of the mechanism by which each hair cell’s activity is tuned to counteract friction. At positions where the hair cells observe a root-mean-square height below the threshold h0, they slowly increase their activity, reducing the effective friction. Otherwise, as in equation relating height and pressure [1, 9, 15], the central part of this sketch, they reduce their activity.

### Resonance from passive impedance

To relate the height and pressure in Eq. 3, we need a mechanical model of the basilar membrane and its surrounding fluid. As is commonly assumed, we take the relationship to be local in space, where it can be written in the frequency domain via the acoustic impedance [11]:

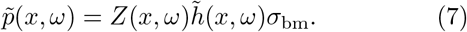

Here, we separate the impedance *Z*(*x, ω*) = *Z*_hc_(*x, ω*) + *Z*_pas_(*x, ω*) into an active component due to hair cells, and the commonly used passive components due to stiffness, inertia, and friction:

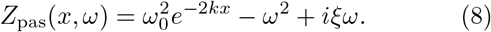

The stiffness decays exponentially in space [16, 17], with *ω*_0_ denoting the resonant frequency at the base of the cochlea. *σ*_bm_ is the mass per area of the BM, and *ξ* the friction per unit mass, which includes viscous losses. A tilde represents a temporal Fourier transform 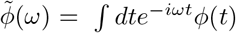. Table I lists all the constants in this model.

**Table I.** Numerical parameters of the human cochlea used in Eq. 1-8. All are taken from Ref. [15].

| Symbol | Value | Description |
| --- | --- | --- |
| $\rho$ | $10^3 \text{ kg/m}^3$ | Density of water |
| $A_{cs}$ | $1.1 \times 10^{-6} \text{ m}^2$ | Average cross-sectional area of a cochlear partition |
| $A_{ow}$ | $3.2 \times 10^{-6} \text{ m}^2$ | Area of oval window (OW) |
| $W_{bm}$ | $2.9 \times 10^{-4} \text{ m}$ | Average width of the BM |
| $\xi_{ow}$ | $500 \text{ s}^{-1}$ | OW dampening constant |
| $\xi$ | $100 \text{ s}^{-1}$ | BM dampening constant |
| $\omega_{ow}$ | $2\pi \times 1500 \text{ rad/s}$ | OW resonant frequency |
| $\omega_0$ | $2\pi \times 20800 \text{ rad/s}$ | Highest resonant frequency of the BM |
| $k$ | $1.38 \times 10^2 \text{ m}^{-1}$ | Stiffness decay coefficient |
| $\sigma_{ow}$ | $18.5 \text{ kg/m}^2$ | Effective areal density of the OW |
| $\sigma_{bm}$ | $5.5 \times 10^{-2} \text{ kg/m}^2$ | Areal density of the BM |
| $G_{me}$ | 21.4 | Gain of the middle ear |
| $L$ | $3.5 \times 10^{-2} \text{ m}$ | Length of the BM |

The passive components of BM impedance lead to the most important feature of cochlear mechanics: spatial frequency discrimination. Resonances occur when the two real contributions to *Z*_pas_ cancel, which happens at a position-dependent resonant frequency

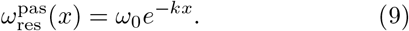

There *Z*_pas_ is purely imaginary. If it were zero, a small change in pressure would result in an infinite change in height, and thus *Z*(*x, ω*) = 0 is a critical point. However, in the real cochlea, a non-zero imaginary part due to friction limits the amplitude of near-resonant displacements. And while the imaginary part of the passive impedance is typically two orders of magnitude smaller than both contributions to the real part [15], it does become significant near the resonant frequency [18, 19]. Since no passive properties can cancel it, hair cells must exert active forces to oppose friction and achieve higher sensitivity.

### Active hair cell contributions

We write the active contribution from hair cells to Eq. 7 in terms of a linear response kernel *g* and a strength *C*:

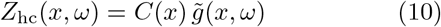

which contributes a pressure *p*_hc_ best understood in the time domain:

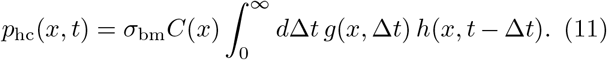

The response kernel *g*(*x*, Δ*t*) characterizes the active force’s dependence on past displacement, and *C*(*x*) controls the strength of the hair-cell force. We initially take *g*(*x*, Δ*t*) ∝ *e*^−*r*(*x*)Δ*t*^, indicating that the hair cell integrates height for a time of order 1*/r*(*x*). This dependence could model, for instance, the concentration of calcium ions which enter when hair cells are displaced and are pumped out at rate *r*, with the accumulated concentration inside the cell controlling molecular motors [8]. To function at both high and low frequencies, we might expect the timescale to vary like 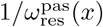 along the BM, and we should certainly expect that different molecular mechanisms will be employed to respond at 200Hz vs 20kHz. Hair cell activity can perturb the resonant frequency away from 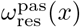, and in general we define *ω*_res_(*x*) by

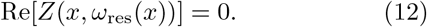

Adjusting the strength of hair cell activity *C*(*x*) is not sufficient to cancel friction for all *x* and *ω*, as the frequency dependence of *Z*_hc_(*x, ω*) comes from 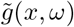 which does not in general match the linear passive term, *iξω*. But adjusting *C*(*x*) does allow us to set *Z*(*x, ω*_res_(*x*)) = 0, cancelling friction along a line in the position-frequency plane. With some important caveats discussed below, we will show that this is sufficient to make the cochlea highly sensitive. To investigate the resulting properties, we assume for now that a fixed fraction *f* (usually 99%) of the passive friction is cancelled at the resonant frequency:

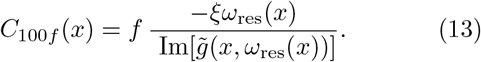

However, cancellation of a large fraction like *f* = 0.99 requires extreme fine-tuning, as a 2% increase in *f* would make the system unstable everywhere. Instead of fine-tuning *C*(*x*) directly, we will later show how hair cells can use local information to robustly tune *C*(*x*) to bring the cochlea to the edge of instability without fine-tuning. But first, we will discuss the qualitative behaviour of the model at *C*_99_(*x*).

### Mode structure of the cochlea

We took a numerical approach to better understand the features of our model, discretizing the BM into *N* units located at *x*_*n*_ = *Ln/N*. Eq. 3 can then be written as ∂_*t*_*X* = *ĴX*, where the state vector *X* concatenates *d*_ow_(*t*), *h*(*x*_*n*_, *t*) for all *n*, their time-derivatives, and any additional entries needed to describe active processes (such as *p*_hc_(*x*_*n*_, *t*) for the one-exponential kernel — see Appendix 1 for details). Diagonalizing the Jacobian *Ĵ* [20] yields modes as eigenvectors with corresponding eigenvalues *λ*_*j*_.

We find, perhaps surprisingly, that the eigenmodes fall into two qualitatively distinct classes, which we term *localized* and *extended* modes, whose eigenvalues and eigen-vectors are plotted in Fig. 2. Each localized mode is strongly peaked at a specific location within the cochlea (orange and yellow in Fig. 2B), and the location *x*_res_ *< L* of this peak is determined by its frequency *ω*_*j*_ = Im[*λ*_*j*_]. But there are a few additional eigenvalues with *ω*_*j*_ *< ω*_res_(*L*) (blue points in Fig. 2A), which correspond to spatially extended eigenmodes (pink and green in Fig. 2B). Increasing the discretization scale *N* increases the number of localized modes, without much effect on either the number of extended modes present (12 in the figure), or their frequencies.

**Figure 2.**
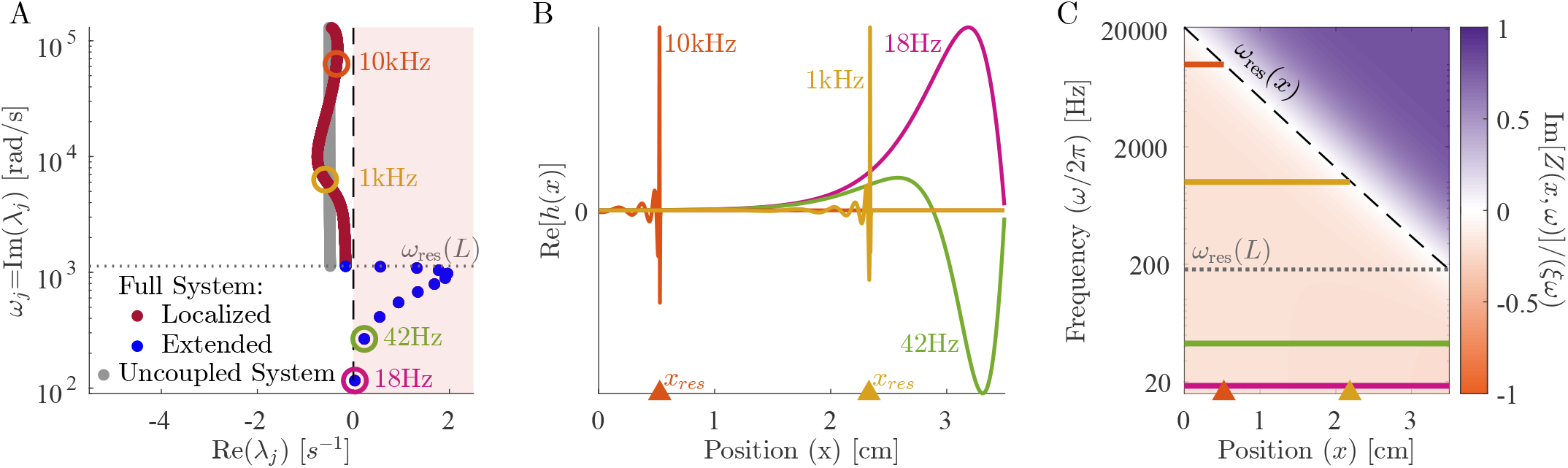
The cochlea exhibits a near-continuum of *localized* modes plus a discrete set of *extended* low-frequency modes. (A) The eigenvalue structure of oscillating cochlear modes. An eigenvalue’s imaginary part determines the oscillation frequency, and the real part determines stability. We define the localized modes (red) as the ones which have a resonant position within the cochlea, i.e., those for which |Im(*λ*_*j*_)| *> ω*_res_(*L*). Extended modes (blue) are those with |Im(*λ*_*j*_)| *< ω*_res_(*L*), so their resonant position would be past the end of the cochlea; e.g., the 18 Hz mode would be resonant at a position of *x* = 5.1 cm. There are 12 extended modes, and *N* − 12 localized modes, approaching a continuum of resonant frequencies at large *N*. The uncoupled system (gray) displays the eigenvalues of *N* independent harmonic oscillators with stiffness, mass, friction, and active force identical to each BM segment (i.e. the roots of *Z*(*x*_*n*_, *iλ*) = 0). (B) Eigenvectors corresponding to the circled eigenvalues. We show the localized modes for 1000 Hz and 10000 Hz and the two lowest-frequency extended modes. (C) Colour map of normalized Im[*Z*(*x, ω*)], the effective net friction, across frequencies and position. On the left of resonance (dashed black line), active processes add more energy than passive friction removes, leading to a negative effective friction (orange). All plots have an active hair cell response kernel *g* ∝ *e*^−*r*Δ*t*^ from Eq. 15 with *α* = 2, *C*_99_(*x*) from Eq. 13, and *N* = 1000. Fig. 3 repeats panels A and C for other choices of response kernel *g*.

The localized modes have been studied in detail [1, 15, 21], and their sharpness in frequency and space is responsible for the remarkable precision with which we can sense pitch. Numerically, we find that the *j*^*th*^ mode is peaked near the resonant position of its eigen-value, *x*_res_ ≈log(*ω*_0_*/ω*_*j*_)*/k* where *ω*_*j*_ = |Im *λ*_*j*_|. In the widely used and qualitatively accurate WKB approximation, the sharp peak at a given frequency is *h*(*x, t*) ∼|*Z*(*x, ω*_*j*_)|^−3*/*4^. Waves approaching from the oval window (*x* = 0) have a decreasing wave speed as they travel right. They slow to zero at *x*_res_, and deposit most of their energy in a so-called critical layer, leaving only an evanescent wave to the right of resonance [10]. Therefore, the stability of these modes is essentially determined by the stability of the local oscillator at resonance. So long as Im[*Z*(*x*_res_, *ω*_*j*_])] *>* 0, active processes are adding in less energy than friction is removing, and these modes are stable, Re(*λ*_*j*_) *<* 0. In the limit of Im[*Z*(*x*_res_, *ω*_*j*_)] → 0, these modes become infinitely peaked, and they can be thought of as essentially uncoupled oscillators acting independently, whose eigenvalues are the roots of *Z*(*x*_*n*_, *iλ*) = 0 (grey points in Fig. 2A). We anticipate that these modes can thus be tuned independently and brought to the edge of instability by choosing *C*(*x*) ≈ *C*_100_(*x*).

By contrast, the extended modes we find are inherently collective. They are defined by having frequencies below the lowest resonant frequency of the cochlea, |Im(*λ*_*j*_) | *< ω*_res_(*L*). These waves travel down the entire cochlea with no evanescent region and can be thought of as sums of right- and left-moving waves that reflect off of the boundary conditions at *x* = 0, *L*. As with a more traditional wave equation, there is a single standing wave mode with no zero crossings (pink in Fig. 2B), one with a single zero crossing (green in Fig. 2B) and so on, with each successive mode having one more zero crossing and a higher characteristic frequency. However, this pattern is cut off at a small number of crossings, corresponding to the maximum frequency still below the resonance of the helicotrema, *ω*_res_(*L*). The number of extended modes, twelve for the parameters used here, thus corresponds to the number of zero crossings of the lowest-frequency localized mode. This number is set by the boundaries of the wave equation, and is independent of the discretization scale except when *N* is very small. While the existence of these extended modes is not a product of active processes, the active processes do influence their eigenvalues and stability.

### Stability of extended modes

While the stability of localized modes is determined by the local balance of active processes and friction at resonance, the stability of extended modes is determined by a combination of these forces along the BM and the dissipation of wave energy out through the oval window. Figure 2C shows the relative net friction

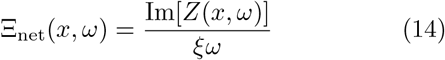

as a function of frequency and position. We observe that for frequencies below the resonant frequency of the helicotrema, *ω < ω*_res_(*L*), it is negative everywhere. As a result, energy is added to the extended modes over the entire BM, leading to their instability. This led us to ask: Are there response kernels for hair cell activity that lead to stable extended modes? And how does requiring the extended modes to be stable limit the response kernels that hair cells might employ?

We propose a simple analytic condition that predicts whether the response kernel will destabilize the extended modes. Fixing some *x* = *x*_0_, we ask whether there is any frequency 0 ≤ *ω < ω*_res_(*x*_0_) at which Im[*Z*(*x*_0_, *ω*)] < 0. In that case, extended modes will experience some negative friction and may be unstable. Even though this simple criterion (shading in Fig. 3B) can be calculated without knowing the height eigenvectors or the coupling to the oval window, it predicts the stability of the extended modes (dots in Fig. 3B) well. (See Appendix 6 for discussion of the oval window’s small effects.) In Fig. 3 we use this criterion, together with the stability obtained from calculating the eigenvalues via the Jacobian, to compare three different response kernels.

**Figure 3.**
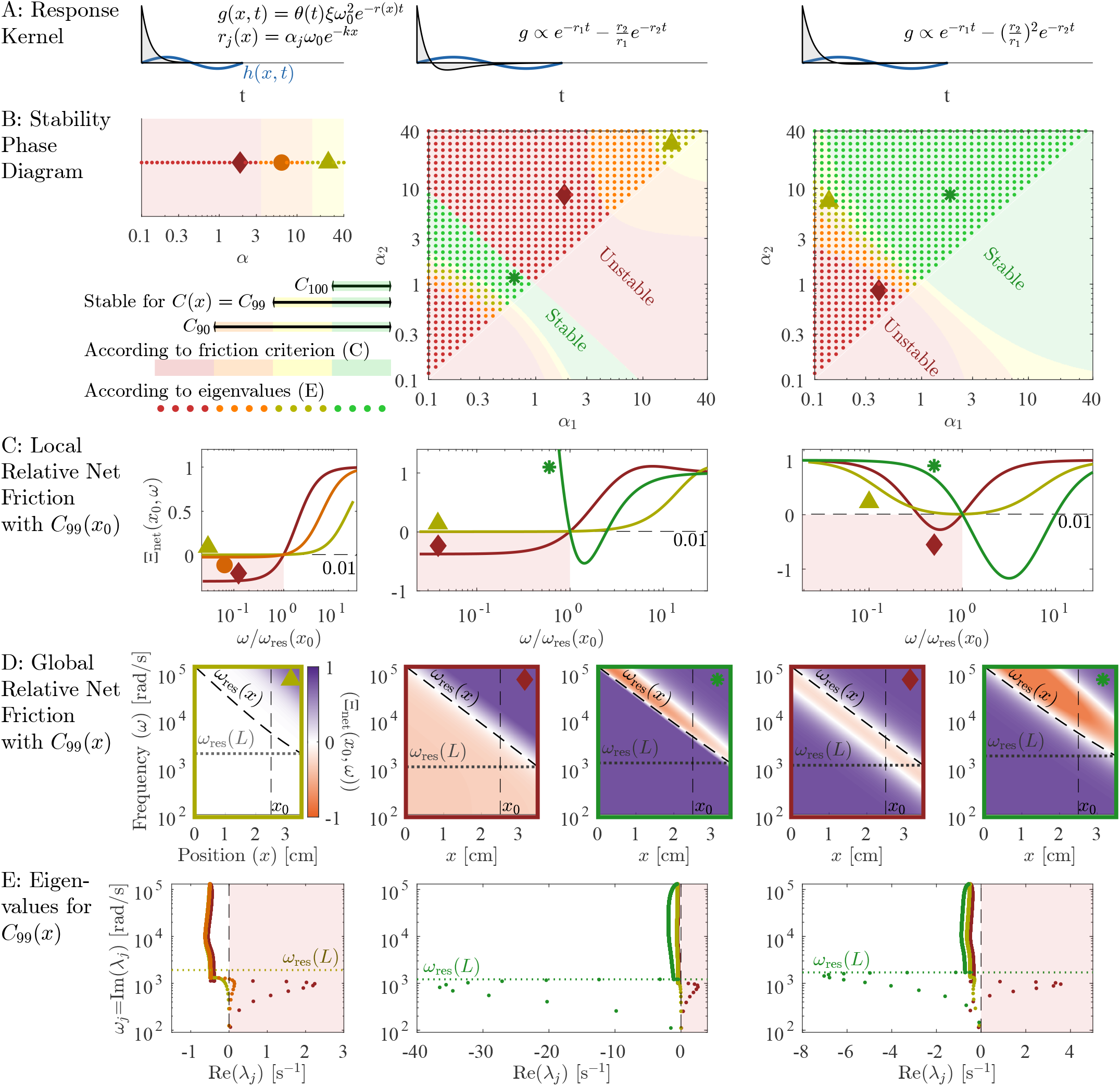
Extended mode stability constrains hair cell response kernels. We consider 3 families of response kernels *g*(*x*, Δ*t*), with the left column showing one exponential with a decay rate *r*(*x*) adjusted by *α*, Eq 15. The middle and right columns have sums of two exponentials whose rates are controlled by *α*_1_, *α*_2_, with two different choices of additive constant, Eqs 16 and 18. (B) Stability phase diagrams, on which green and yellow points indicate parameters *α*_1_, *α*_2_ at which there are no eigenvalues with positive real part when hair cells are tuned to cancel 99% of passive friction on resonance, *C*(*x*) = *C*_99_(*x*) from Eq. 13. If the hair cells are instead tuned to cancel 90% of passive friction (*C*_90_), then more cases become stable, indicated by orange points. If they cancel 100% of passive friction (*C*_100_), then only the green points are stable. Background colour indicates stability according to the criterion of having Im *Z*(*x, ω*) *>* 0 for all *ω < ω*_res_(*x*). Points marked by large symbols are plotted in lower panels, always with *C*_99_(*x*). (C) At a particular position *x*_0_ = 2.5cm, we plot relative net friction Ξ_net_(*x, ω*), for frequencies *ω* above and below the resonance frequency *ω*_res_(*x*_0_). By definition of *C*_99_, this is equal to 1 − 0.99 at *ω* = *ω*_res_(*x*_0_). The friction criterion states that any negative values at *ω < ω*_res_(*x*_0_) will produce instability (red background). (D) Relative net friction for all *x* and *ω*. By definition of *C*_99_, this is 0.01 along the resonance line *ω* = *ω*_res_(*x*) (dashed). Positive values (purple) indicate energy loss, while negative values (orange) indicate energy gain, potentially leading to instability. (E) Mode eigenvalues, for the same selected parameters *α*_1_, *α*_2_ indicated by large symbols on the phase diagrams. Any eigenvalue with Re(*λ*_*j*_) *>* 0 is an unstable mode (red background). Eigenvalues in (B,E) were calculated at *N* = 1000.

### Single exponential kernel

So far we have used the response kernel introduced below Eq. 11, which integrates height for a time we assumed to be similar to the period of a wave resonant at that location:

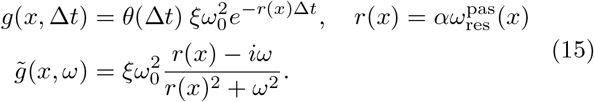

Here *θ*(Δ*t*) is the Heaviside function that ensures a causal response. As we have seen in Fig. 2, for *C*(*x*) = *C*_99_(*x*) and *α* = 2 this choice leads to unstable extended modes since the net friction is negative everywhere for low frequencies. At the same time, the net friction is, by construction, slightly positive at the resonant frequency.

In Fig. 3C left we plot the relative net friction at a particular location *x*_0_ for several values of *α*. This Ξ_net_(*x*_0_, *ω*) is monotonically increasing as a function of frequency and is given by 1 − *f* at resonance, by definition. Thus, as *f* approaches 1, net friction always becomes negative at lower frequencies, and hence our criterion predicts instability. In general, this criterion predicts stability only for 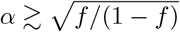, in approximate agreement with our eigenvalue results (dots in Fig. 3B). Consequently, exponential kernels cannot bring localized modes to the edge of instability (*f*≈1) without destabilizing extended modes, and thus we do not expect to find active processes with this form in hair cells. Can other response kernels stabilize these extended modes?

### Approximate derivative kernel

The most common approach when studying cochlear dynamics is to assume friction is uniformly cancelled [9, 15, 21], equivalent to taking a response kernel that implements an instantaneous derivative *p*_hc_(*x, t*) ∝ ∂_*t*_*h*(*x, t*) or 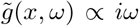. So if *f* ≤ 1, all modes at every position and frequency would experience a positive or zero relative net friction leading to stable extended modes. However, an instantaneous derivative requires an infinitely fast response of hair cells. With a finite response time, one can approximate a derivative by a sum of two exponentials:

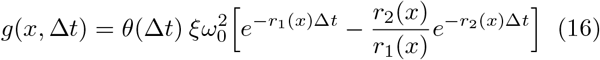

with 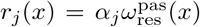. The relative coefficient −*r*_2_*/r*_1_ between the two terms ensures that ∫*d*Δ*t g*(*x*, Δ*t*) = 0, so that a time-independent shift in *h* has no effect on the response. In the limit *α*_1_, *α*_2_→ ∞, this response kernel approaches an instantaneous derivative.

Figure 3B middle shows the stability phase diagram for this response kernel as a function of *α*_1_ and *α*_2_. For *f <* 1, the extended modes are stabilized for large *α* as in the case of a single exponential. But there is also a narrow range with *α*_1_*α*_2_ ≲ 1 (green in Fig. 3B) where the extended modes can be stable all the way to *f* = 1. This true stability occurs because relative net friction has a negative slope at the resonance (green in Fig. 3C), so that negative net friction occurs at frequencies higher than resonance (orange in Fig. 3D). Perhaps surprisingly, this region of true stability occurs for low rates, *α*_1_*α*_2_ ≲ 1. Hence, hair cells could implement kernels with this form, but not in the regime where they closely approximate an instantaneous derivative.

### Zero-derivative kernel

Based on our simple condition for stability, can we construct a response kernel that leads to stable extended modes for a broader range of parameters? Since the simple analytic condition focuses on the behaviour of Ξ_net_(*x, ω*) at low frequencies, we approximate it by a Taylor series in *ω*:

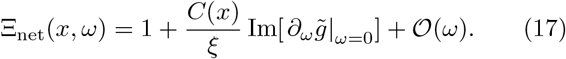

A simple condition for positive net friction for small *ω* independent of *C*(*x*) is thus given by 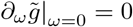, which in the time domain reads 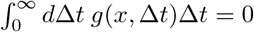. This condition can be met by a sum of two exponentials weighted as follows:

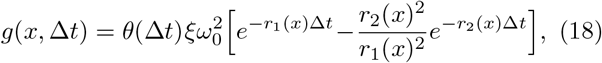

where 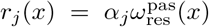 as before. With this kernel, Fig. 3 right, there are three possible cases: In the first case, friction is positive everywhere. This case is only possible for *f <* 1 and gives rise to the yellow and orange area in the stability phase diagram. In the second case, friction is negative in a band of frequencies below the resonance frequency, leading to instability of the extended modes, see red area in Fig. 3B right. Finally, friction is positive below the resonant frequency but negative for a band of frequencies above the resonance frequency (green area). Importantly, this stable regime occurs for points *α*_1_*α*_2_ ≳ 1 and therefore does not require much tuning of parameters.

Taken together, our results suggest that the extended modes are unstable if the net friction is negative for (a band of) frequencies lower than the resonance frequency. In principle, there are two ways to avoid this situation: either net friction is positive everywhere, or it is negative for a band of frequencies larger than the resonance frequency. The former can only occur if the net friction at resonance isn’t fully cancelled (*f <* 1), thus making this regime less useful for tuning the localized modes to the edge of instability. The latter regime also works for fully cancelled friction *f* = 1. It only occurs in a narrow parameter regime if the kernel approximates a derivative (Fig. 3 middle) but can be greatly enhanced if the kernel 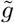 instead exhibits a zero first derivative at *ω* = 0 (Fig. 3 right). It is worth noting that for all forms of the kernel 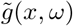 considered here, localized modes are at least marginally stable for all *f* ≤ 1.

### Independent tuning of localized modes

Having established conditions on the stability of extended modes, we now turn to how hair cells can tune all localized modes to the edge of instability. Since the friction term in *Z*_pas_ is small compared to the canceling real parts, the relatively small absolute contribution of *Z*_hc_ will dramatically affect mechanics only where the real part of *Z* is near 0. As a result, for kernels with stable extended modes, we propose that hair cells only need to cancel friction for the localized mode peaking at their location. Fig. 4 shows that if we reduce friction to near zero at a specific location, we only see a noticeable amplification of frequency modes that peak near that position (Fig. 4C). Off-resonance amplification (Fig. 4B) looks qualitatively identical to a passive system (Fig. 4A). Tuning the cochlea therefore only requires that friction is cancelled very near the resonant frequency (Im[*Z*(*x, ω*_res_(*x*))] = 0 for all *x*), a far less stringent limitation than globally nullifying friction (Im[*Z*(*x, ω*)] = 0 for all *x, ω*).

**Figure 4.**
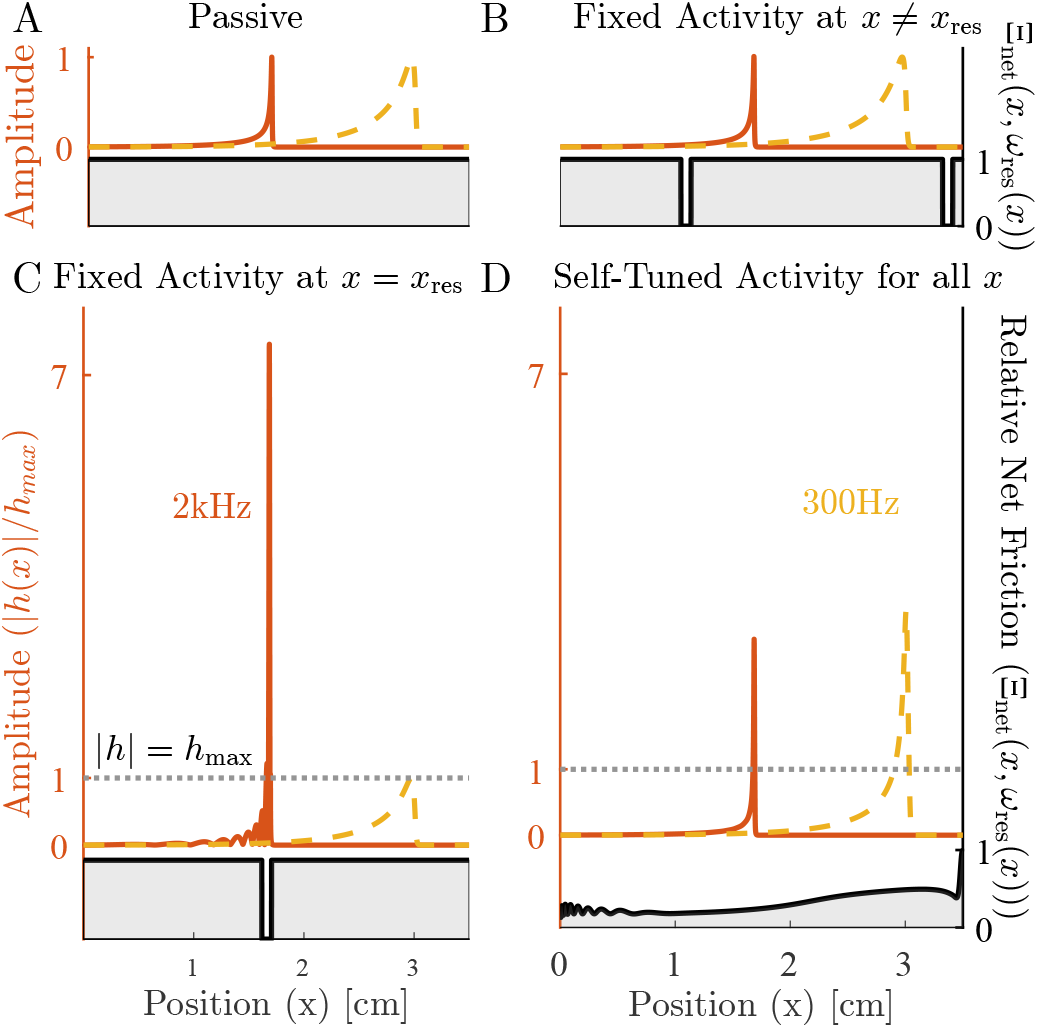
Canceling friction has a predominantly local effect. We show the amplitude of the cochlea’s response to pure tones at 2000 Hz (orange) and 300 Hz (yellow) for (A) a passive cochlea, *C*(*x*) = 0, which by definition has Ξ_net_(*x, ω*) = 1, (B) friction reduced off-resonance, (C) friction reduced at the resonant position for 2000 Hz, giving an over 7-fold amplification, and (D) a cochlea self-tuned using Eq. 19. The left axis shows the amplitude of the response, normalized to peak at 1 in the passive case. The right axis shows the net effective friction at resonance Ξ_net_(*x, ω*_res_(*x*)), Eq. 14, shown in gray in all panels.

### Self-tuning of active process strength

So far, we have shown that requiring stability of extended modes puts constraints on viable active hair cell responses and that for stable kernel choices all localized modes can be brought to the edge of instability simultaneously. In part this tuning is possible because the effects of *C*(*x*) are local both in space and in frequency. However, for strong amplification, it also requires the local hair cell activity strengths *C*(*x*) to be tuned very close to *C*_100_(*x*), Fig. 5. So, how can hair cells find the region where the net friction is cancelled (almost) perfectly along the line of resonant frequencies and thereby bring the set of localized modes to the proximity of their individual instabilities? Inspired by how isolated hair cells in non-mammalian vertebrates as, for example, bullfrogs can tune themselves to their Hopf bifurcation [8], cells could take advantage of the extreme sensitivity of the membrane displacement amplitude near *C*_100_(*x*) to indirectly tune to this region, by measuring the amplitude of local displacements. This indirect tuning works much more robustly than tuning *C*(*x*) directly because the size of local oscillations in BM height are an order parameter for a dynamical bifurcation where *C*(*x*) is a control parameter.

**Figure 5.**
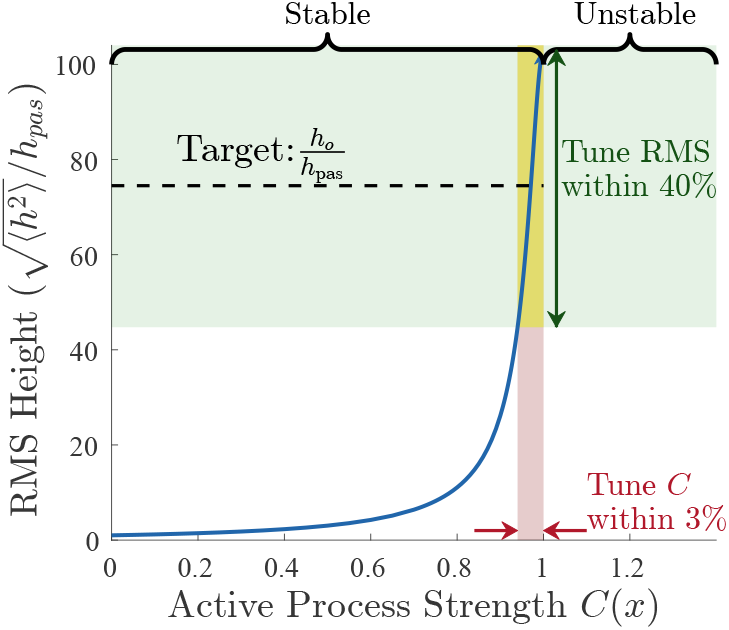
RMS height diverges as active process strength cancels passive friction. At a fixed position we plot RMS height as *C*(*x*) is varied, using the instantaneous derivative kernel 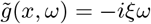 for which friction is fully cancelled when *C*(*x*) = 1. We show how a 3% change in *C*(*x*) corresponds to a much larger change in RMS heights. Thus, controlling *C*(*x*) directly would require fine-tuning (red-shaded area), but feedback based on the height would only require *h*_0_ to fall on the steep part of this curve (green-shaded area). The region to the right of *C*(*x*) = 1 is unstable.

We therefore consider feedback which, rather than directly implementing *C*(*x*) ≈ *C*_100_(*x*), instead adjusts *C*(*x*) to target an order parameter, the RMS displacement: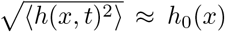. Because the effect of *C*(*x*) is predominantly local in *x* (Fig. 4), this can be implemented by local feedback. Thus we consider adding additional slow dynamics to the model:

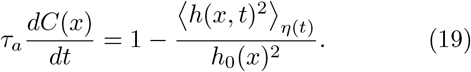

Here we assume that the timescale of the feedback *τ*_*a*_ is much longer than the longest timescale of the sound-driven dynamics: *τ*_*a*_ ≫1/20Hz. And we imagine that the system is externally driven by input from the ear canal, and model incoming sound as uncorrelated Gaussian noise in pressure: *P*_ec_(*t*) = *η*(*t*). The temporal average needed for RMS displacement has thus been replaced by an average over this noise process — see Appendix 2 for details. (Due to symmetry, the average of *h*(*x, t*) vanishes, ⟨*h*(*x, t*)⟩_*η*(*t*)_ = 0, thus feedback from the height squared is the first nonzero moment.) In plots below we use as the target *h*_0_(*x*) five times the passive RMS displacement.

We find that this scheme can bring each mode to the edge of instability without fine-tuning any fixed parameters (green in Fig. 5). *C*(*x*) is a dynamical function approaching a steady-state value that only weakly depends on *h*_0_. This behaviour is reminiscent of systems that exhibit self-organized criticality [8, 13, 14, 22], where a slowly varying control parameter is tuned via feedback from a fast order parameter.

### Robustness to perturbations

This section examines two possible sources of variability and how, despite them, the system can still find its critical point. First, we consider a case in which the middle 10% of the cochlea is forced to have zero activity. We observe that points away from this region still reach their critical points and that inactive points very close to the edges of inactivity can be partially amplified (Fig. 6A). This observation reinforces the local nature of these active processes, showing that only friction at a given position needs to be reduced to achieve amplification for that position’s resonant frequency.

**Figure 6.**
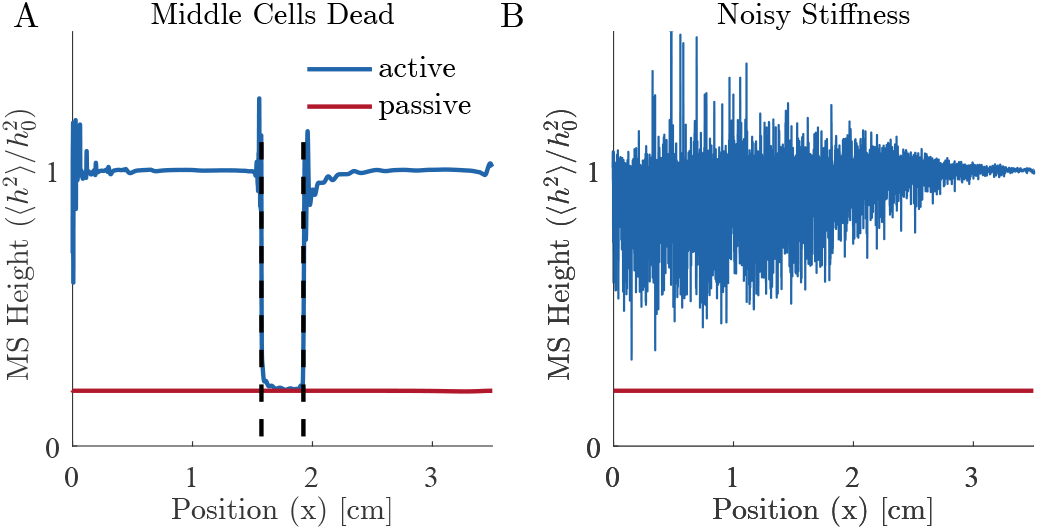
The cochlear self-tuning is robust to perturbations. Mean-square height for (A) a simulation in which the middle 10% of cells are inactive, i.e. *C*(*x*) = 0 for 0.45*L < x <* 0.55*L*, (B) a simulation for which the exponentially decaying stiffness 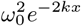 has multiplicative white noise with a standard deviation of 1%. Note that we use a discretization scale of *N* = 4000, which might not be large enough to resolve the peaks of the high-frequency localized modes close to *x* = 0 and lead to the observed spikes there in (A).

Another important part of our model is the stiffness of the basilar membrane, whose position dependence determines the resonance position of the different frequencies. To check how robust self-tuning is to slight changes in the stiffness profile, we consider a system in which the stiffness of the BM is noisy, replacing 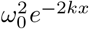 with 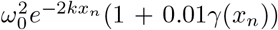, where *γ*(*x*_*n*_) is iid standard Gaussian noise. Figure 6B shows the resulting steady-state mean-square height profile. We see that self-tuning still achieves similar average enhancement, demonstrating that the exact stiffness profile is not necessary for amplification in the cochlea. However, the resulting system is quite noisy. We believe that this noisiness is due to the fact that the stiffness no longer decreases monotonically everywhere. If, as a result, the real part of the impedance is negative already before resonance, the travelling wave is exponentially suppressed and decays rapidly there. In order to match the target RMS height, hair cells at this location try to overcompensate by considerably increasing their activity. The resulting increase in *C*(*x*), however, also affects neighboring points and leads to the large spikes seen in Fig. 6B.

## Discussion

The high sensitivity, dynamic range, and frequency resolution of human hearing are all thought to arise due to proximity of individual oscillators to Hopf bifurcations, driven by hair cell activity that effectively reduces friction [21]. Here we introduce a wave equation for the basilar membrane that includes hair cell activity in terms of a generic response kernel, with a position-dependent activity strength, and analyze its mode structure.

We find modes that peak at a resonant position, which we call localized modes, and argue that it is these modes whose Hopf bifurcations enable the fidelity of hearing. Although different spatial locations are, in principle, coupled by fluid flow and influenced by hair cell activity throughout the cochlea, we demonstrate that for small friction, the localized modes become so sharply peaked that different positions are nearly uncoupled. In this small friction limit the amplification of each mode is dependent only on the activity of hair cells near the resonant position. Thus, a simple, local feedback scheme for hair cell activity strength can tune all localized modes to the edge of instability.

Surprisingly, however, we also find a second set of modes, which we call extended modes. These are standing-wave-like, and couple to essentially all hair cells. We find that requiring stability of extended modes provides strong limitations on viable hair cell responses. In particular, our results suggest that approximations to the derivative kernel, which is often implicitly assumed to underly hair cell activity [9, 23], generically lead to unstable extended modes. Other kernels might be better suited, and we find criteria for the temporal shape of active processes which hair cells must obey. An interesting question for further research is to understand which proposed molecular mechanisms satisfy these criteria. For kernels which obey these criteria the localized modes become unstable at smaller activity strength than the extended modes do, and so local tuning can bring the localized modes independently to the edge of instability. In the self-organized steady-state of our model, effective friction is only cancelled along the one-dimensional resonant line through the two-dimensional space of frequency and cochlear position. This is in contrast to existing models that analyze a nonlinear cochlea by assuming that friction has been set globally to (near) zero [1, 9, 11].

Models for isolated hair cells without a cochlea use a feedback scheme similar to ours, with a control parameter, usually calcium activity, tuned towards a Hopf bifurcation [6, 8]. In both cases, tuning is based on the (local) order parameter, here the BM displacement, and works robustly due to the large susceptibility at the critical point. Using criticality for sensing might indeed be common in biological systems. For instance, for E. coli chemosensing, we and others have proposed that cell receptor arrays tune themselves close to criticality to detect small changes in concentration [24, 25]. In the neural realm, we have suggested that proximity to a bifurcation of the voltage dynamics might underly the incredible temperature sensitivity of pit vipers [26] and allow fruit flies to reliably extract odor timing information for navigation [27]. Furthermore, it is thought that the schooling behaviour of fish, flocking of birds and swarming of insects are near a phase transition to optimize collective computation [28–30]. Finally, it has been shown that an anti-Hebbian learning rule in neurons whose connectivity is suppressed in response to activity can lead to a self-organized dynamical critical state [31–33].

Whether the order-parameter based feedback scheme presented here is implemented in hair cells could be tested by measuring the cochlea’s response when given a continuous signal at a limited frequency bandwidth. We expect hair cell activity at positions with a resonant frequency close to that of the signal to decrease while having a minimal effect for positions further away. Directly testing this prediction would require measuring the response in a live cochlea, a difficult but perhaps feasible task with emerging optical coherence tomography techniques [34, 35]. It might also be possible to indirectly test this prediction with simpler psychoacoustic methods.

While our model provides a mechanism for the cochlea to tune itself using only local information, there are hints that there is at least some nonlocal feedback. Under contralateral stimulation (sound played in the opposite ear), the frequency of known otoacoustic emissions (OAEs) shifts. This phenomenon is thought to be due to changes in outer hair cell activity induced by signals from neurons in the MOC bundle [36]. In our model, such shifts also occur naturally if contralateral stimulation globally decreases *C*(*x*): If the contribution *Z*_hc_ of hair cells to the impedance is not purely imaginary, a change in *C*(*x*) shifts the resonant frequency of that position (and thereby changes the frequency of otoacoustic emissions) by a small amount 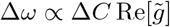. Such frequency shifts have indeed been observed in lizards [37]. Should they also be observed in human hearing when stimulation happens in the same ear as the measurement of OAEs, this could lend further credence to our local order parameter-based feedback.

Early evidence for the importance of a Hopf bifurcation in hearing came from characteristic nonlinearities, including that the BM wave amplitude grows as the 1*/*3 power of sound amplitude and a prominent third harmonic in evoked otoacoustic emissions [1, 9, 21]. Our model is explicitly linear, which we expect to be a good approximation far from the edge of instability. But when hair cell activity is strong enough to precisely bring the impedance at resonance to zero, the small nonlinearities of the basilar membrane become the dominant restoring force. It is thus an interesting question for future research how these nonlinearities interact with extended modes.

In addition to making a linear approximation, we discretized the cochlea to understand its mode structure. This discretization separates the BM into segments of length *L/N* where *N* typically ranged from 1000 to 4000 in our numerics. The real cochlea also contains a small spatial scale, set by the length at which lateral coupling dominates, around 20*μ*m [16, 38], corresponding to around 2000 independent segments. We expect that the number, shapes and eigenvalues of the extended modes will be independent of details at this small spatial scale. However, the details of the short length-scale physics might influence localized modes in interesting ways.

Frequency discrimination and signal amplification in the range of 20 − 1000Hz remains an area of active research [39]. Since the extended modes exhibit resonant frequencies below the lowest resonant frequency of the basilar membrane (165Hz), they could potentially contribute to the cochlea’s low-frequency sensitivity. Indeed, experimental measurements of BM dynamics have revealed a constant phase slope near the cochlear apex, indicating that low-frequency waves reach the helicotrema [40, 41]. This observation aligns with the characteristics of the extended modes we present and the exploration of these extended modes and their impact on hearing continues to be an exciting avenue for future research.

## Acknowledgments

We thank James Hudspeth and the members of the Machta group for helpful discussions, and Pranav Kantroo, Mason Rouches, Derek Sherry and Jose Betancourt for constructive comments on the manuscript. This work was supported by the Deutsche Forschungsgemeinschaft (DFG, German Research Foundation) Projektnummer 494077061 (IRG), and by NIH R35GM138341 (BBM) and a Simons Investigator award (BBM).

## Appendix

In this appendix, we will first demonstrate how we determine the stability of the basilar membrane in a finite element approximation. Then, we will show how we calculate the variance in height and use this to tune *C*(*x*). Next, we will look at the eigenvalue structure, followed by a discussion on how this structure scales with discretization. After this, we consider an alternative response kernel and some perturbations to the existing response kernels. Finally, we look at the effect of increasing oval window friction.

### Appendix 1. Jacobian and Stability

Our passive cochlear model is based on work from Ref. [15], with a few changes in notation and any larger differences discussed as they are introduced. We model the cochlea with two compartments. First we consider the bulk of the cochlea where sound induces flux of water in the upper and lower compartment; we define *j*(*x, t*) = *j*_lower_(*x, t*) − *j*_upper_(*x, t*) as the difference in volume current between the scala tympani and scala vestibule. According to force balance, this creates a corresponding pressure difference *p*(*x, t*) = *p*_lower_(*x, t*) −*p*_upper_(*x, t*), between the two compartments,

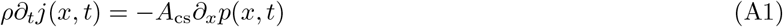

where *A*_cs_ is the average cross-sectional area of a cochlear compartment and *ρ* is the density of water. The fluid flux propagates down the cochlea, creating a displacement *h*(*x, t*) of the BM, which we call height. The fluid flux obeys a continuity equation

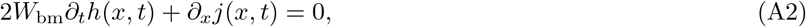

where *W*_bm_ is the width of the BM. We can eliminate *j*(*x, t*) from Eq. A1 and A2 to arrive at a modified wave equation,

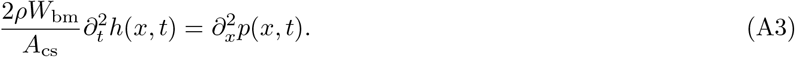

For our numerical solution, we use a finite element approximation to this equation where we break the cochlea into *N* points separated by a distance 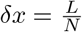, and we label each point *x*_*n*_ = *nδx* with *n* = 1, 2, …, *N*. Now Eq. A3 becomes,

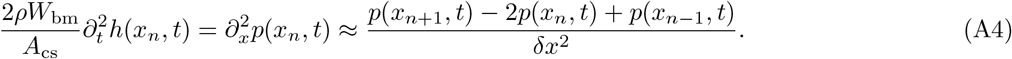

We now turn our attention to the boundary conditions. At the left-hand side, we have a Neumann boundary condition where the lateral displacement of the oval window *d*_ow_(*t*) creates a flux of fluid, which propagates down the cochlea and induces an equal but opposite flux at the round window, due to fluid incompressibility. Via Eq. A1, this flux leads to a pressure gradient:

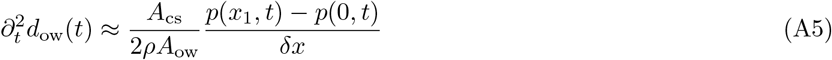

where *A*_ow_ is the area of the oval window. The oval window itself acts as a damped harmonic oscillator:

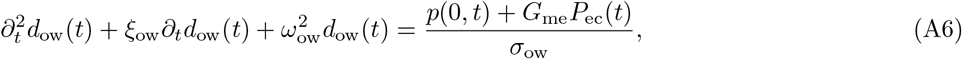

where *ξ*_ow_ is its dampening constant, *ω*_ow_ the middle ear resonance, *P*_ec_(*t*) the pressure in the ear canal, *G*_me_ the gain of the middle ear, and *σ*_ow_ the (2D) areal density of the oval window. At the cochlea’s apical end *x* = *L*, a gap in the basilar membrane (the helicotrema) suggests zero pressure difference via the Dirichlet boundary condition [15]:

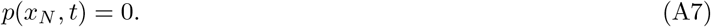

Now assuming that *P*_ec_(*t*) = 0 we can write the discretized system in the form,

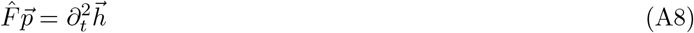

We will introduce the shorthand *p*(*x*_*n*_, *t*) = *p*_*n*_and *h*(*x*_*n*_, *t*) = *h*_*n*_ as we write 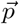 as an *N* column vector,

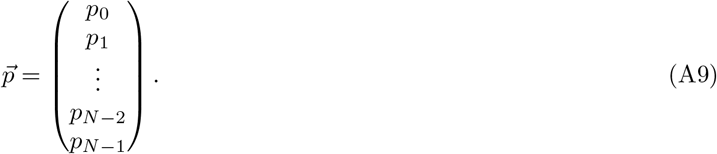

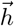 is an *N* vector of BM displacement with *h*(0, *t*) replaced by *d*_ow_(*t*),

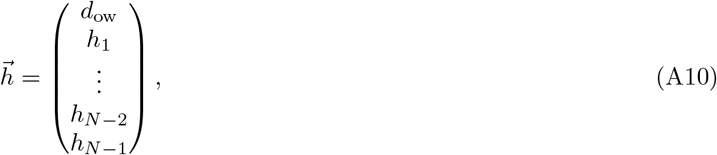

and 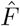 is a modified finite difference matrix,

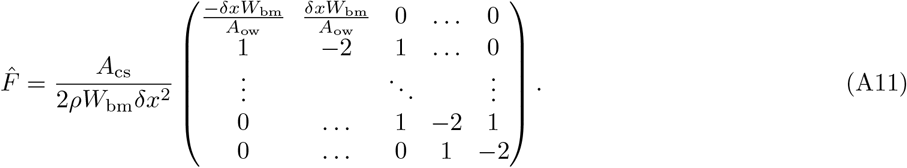

Note that we have used the right-hand side boundary condition (*p*_*N*_ = 0) to eliminate the last row of the matrix.

We use a modified version of the formalism from Ref. [20] to write the dynamics as

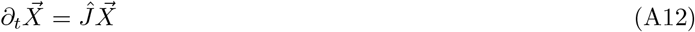

where *Ĵ* is the system’s Jacobian and 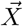 is state vector concatenating *d*_ow_(*t*), ∂_*t*_*d*_ow_(*t*), *h*(*x*_*n*_, *t*), ∂_*t*_*h*(*x*_*n*_, *t*) with *n* = 1, 2, 3…, *N* −1 and any additional degrees of freedom needed to describe active processes. Here, we show this procedure explicitly for *g*(*x, t*Δ) ∝ *e*^−*r*(*x*)Δ*t*^.

To begin with, we write the height-pressure relation as follows,

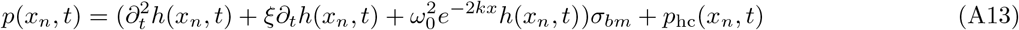

The stiffness 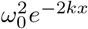 is exponentially decaying in space, with *ω*_0_ denoting the resonant frequency at the base of the cochlea. *σ*_bm_ is the mass per area of the BM, and *ξ* the friction per unit mass. Our main departure from Ref. [15] is that we treat friction along the cochlea as a constant. The active hair cell contribution to the pressure is

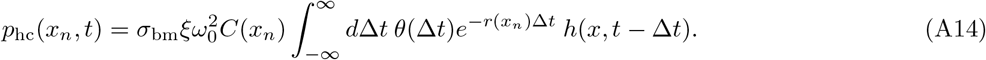

for some positive function *r*(*x*_*n*_) and *C*(*x*_*n*_) defined in the main text. Direct computation of ∂_*t*_*p*_hc_(*x*_*n*_, *t*) reveals,

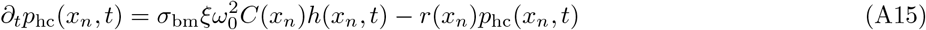

For this particular form of *p*_hc_(*x, t*) we write the vector 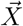 as column vector of length 3*N* − 1 as follows:

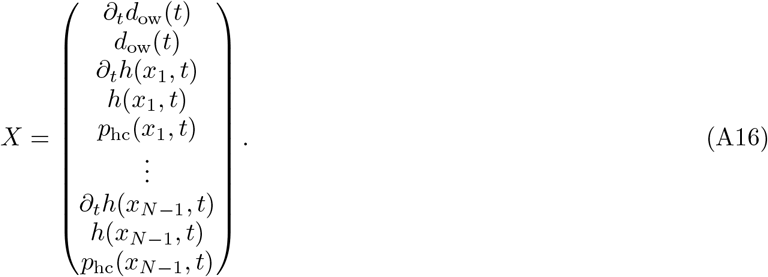

With this choice,

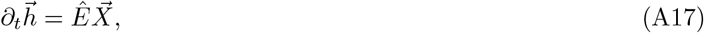

where 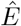 is a *N ×* (3*N* − 1) block diagonal matrix with the following form,

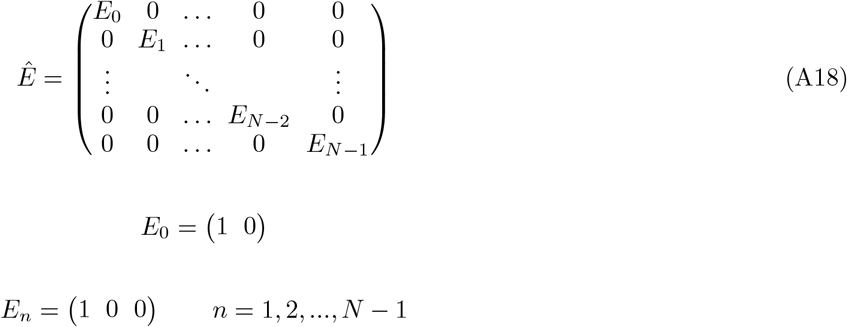

In order to get the Jacobian of 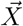 we must first take its time derivative. We can express this time derivative as a sum contribution from 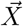, with the prefactor of each term in 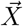 given by a matrix 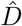, and contribution from the pressure across the BM 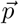, with the prefactor of each term in 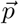 given by a matrix 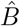,

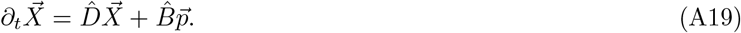

Here 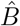 is a (3*N* − 1) *× N* matrix with the following form,

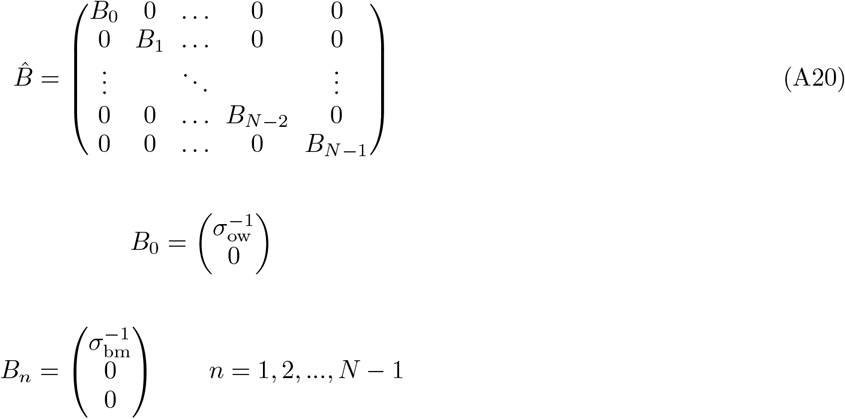

And 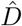 is a (3*N* − 1) *×* (3*N* − 1) matrix with the following form,

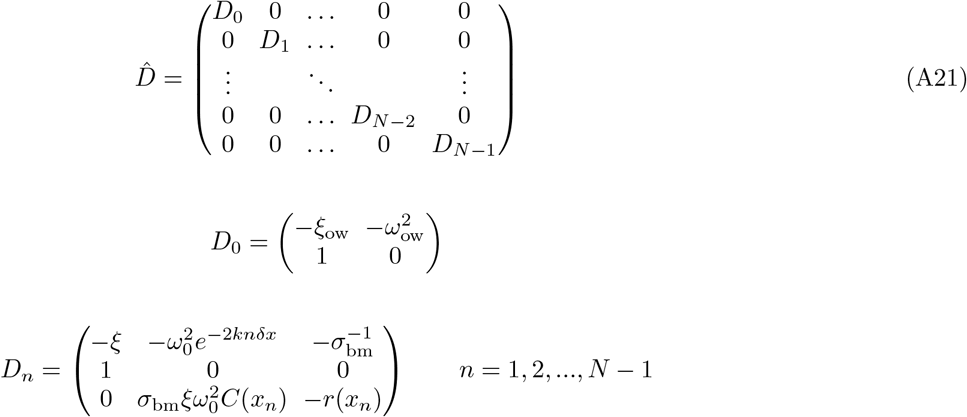

We can then combine Eq. A8 with Eq. A19 to write,

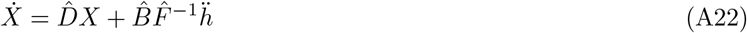

Then substituting equation A17 yields

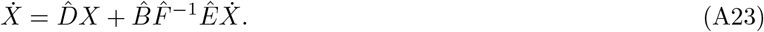

Finally, a simple rearrangement yields,

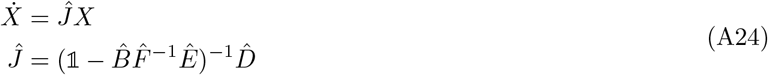

where 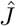 is the Jacobian of the system, a (3*N* − 1) *×* (3*N* − 1) matrix.

### Appendix 2. Variance of Height

In the main text, we introduce a tuning scheme in which the strength of active processes changes with the variance of BM displacement. This section explains how we calculate ⟨*h*^2^⟩. We start by writing Eq. A3 and A13 in Fourier time,

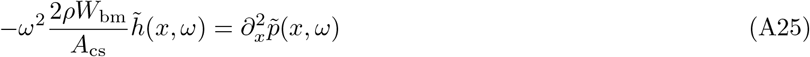

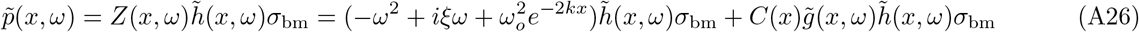

where a tilde represents a temporal Fourier transform, 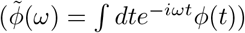. We want to discretize the system into N segments, each segment obeying,

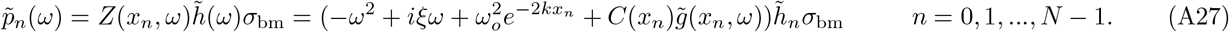

Using a discrete derivative,

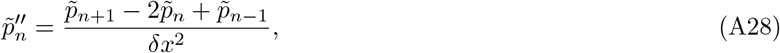

we rewrite the right-hand side boundary conditions in Fourier time,

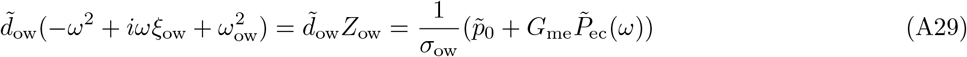

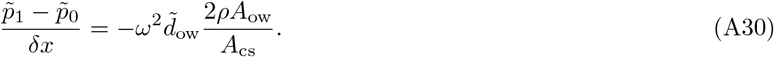

Then we use this equation to define 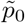 in terms of 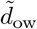 and 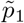 and eliminate 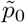 in Eq A29. We assume 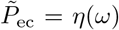 where *η*(*ω*) is Gaussian white noise ⟨*η*(*ω*)*η*(*ω*^*′*^)⟩ = *σ*^2^*δ*_*ω*,−*ω*′_.

It is easiest to work with pressure, so we use Eq. A26 to write height in terms of pressure difference. Then, we can write this system in matrix form.

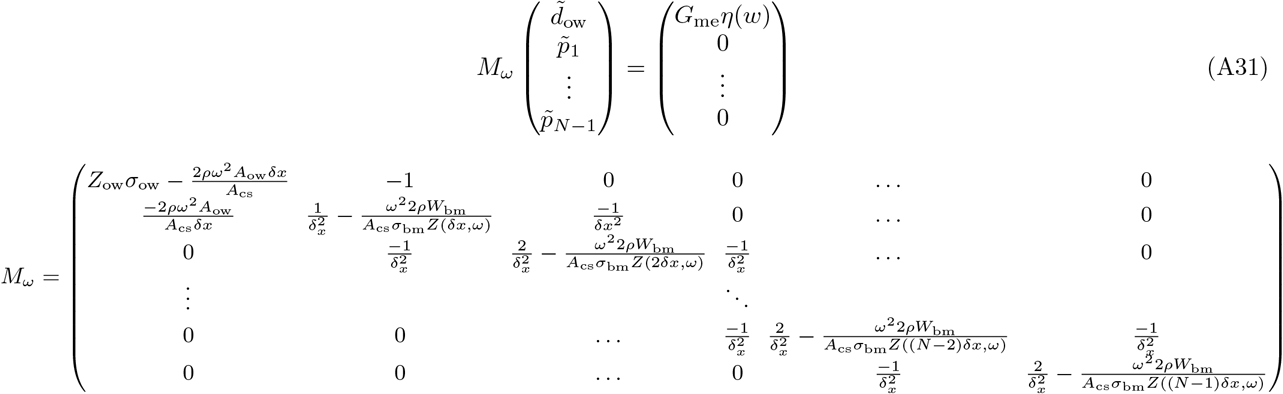

To find the pressure at any given location, we have to invert *M*_*ω*_ (one can make use of the tridiagonal structure of *M*_*ω*_ for a computational speed-up [42]).

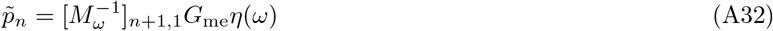

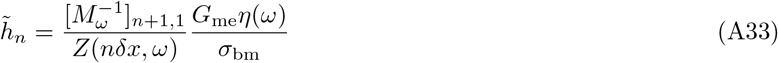

We can then make use of the Wiener-Khinchin theorem, which states,

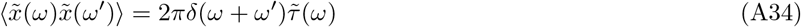

where 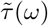 is the Fourier transform of the system’s autocorrelation function. Applying this theorem to our systems yields for the variance of the height

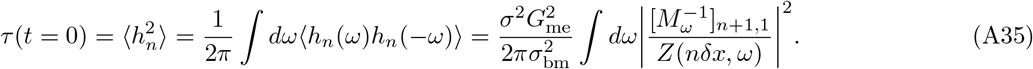

We then use this to iterate feedback on *C*(*x*_*n*_) of the form,

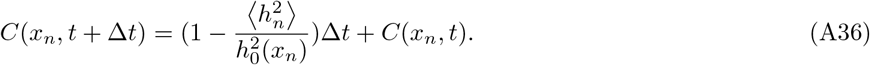

It is worth noting that this feedback assumes an infinite separation in time scale between the average 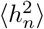 and the feedback on *C*(*x*).

### Appendix 3. Eigenvalue structure

In the case of a single exponential response kernel, the Jacobian has 3*N*−1 eigenvalues. 2(*N*−1) of these come from the position and velocity of the basilar membrane and are oscillating modes. The eigenvectors and corresponding eigenvalues of these modes are the localized and extended modes discussed in the main text. 2 modes correspond to predominately oval window motion. The remaining *N*− 1 modes come from active processes and are non-oscillating (imaginary part of 0). These modes have an eigenvalue with a real part far less than 0; and are absent if active processes are excluded. Fig A1 shows the eigenvalue structure for 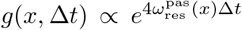. Fig A1A shows the full eigenvalue spectrum, and Fig A1B shows those included in the main text. We exclude eigenvalues with Im *λ*_*j*_ = 0 as these have a real part 4-5 orders of magnitude lower than those of oscillating modes and will quickly decay to 0.

**Figure A1.**
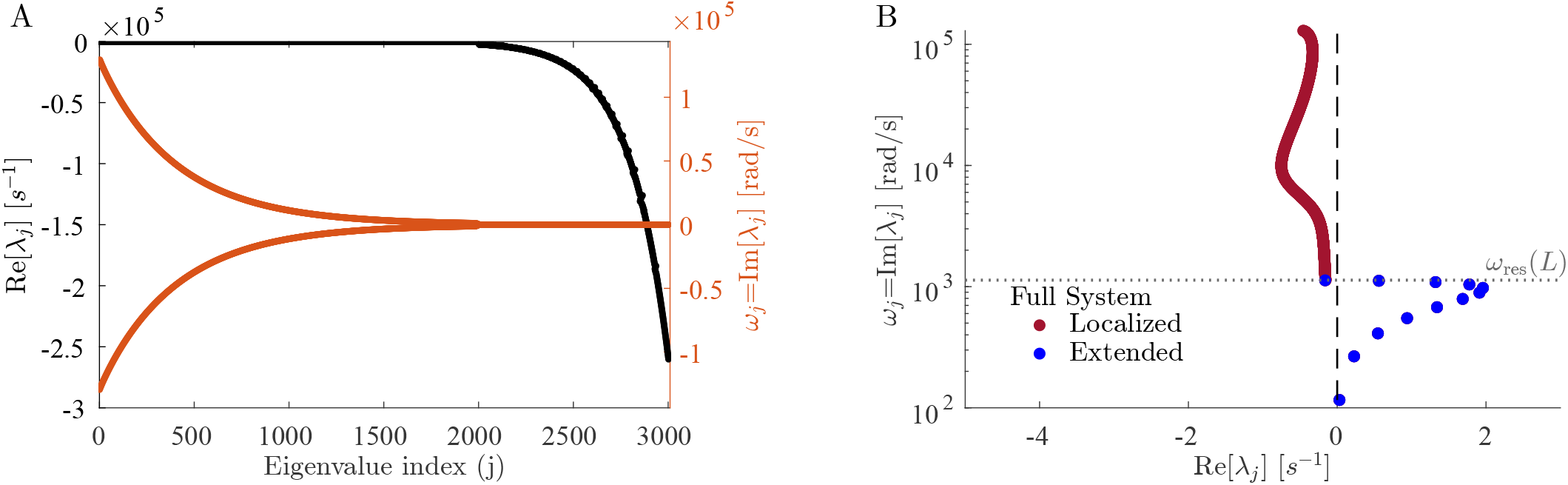
The full eigenvalue structure. A) All eigenvalues are rank-ordered by the magnitude of their complex part. Note that approximately a third of eigenvalues have an imaginary part exactly equal to 0, and there is a large spread in the value of the real part of the eigenvalues. B) Eigenvalues in a semi-log plot as in main text Fig. 2. Note that all eigenvalues with an imaginary part less than or equal to 0 are omitted. However, this plot contains all the details relevant to our purposes for two reasons. First, since all complex eigenvalues come in conjugate pairs, eigenvalues with a negative frequency look exactly like those plotted with Im[*λ*] → −Im[*λ*]. Second, all eigenvalues with Im[*λ*] = 0 are always stable in every simulation we have run. These modes do not oscillate and have a large negative real part; they would fall to 0 very quickly if stimulated and do not affect the behaviour discussed in the main text. For these plots 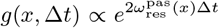 with the friction 99% cancelled (*C*(*x*) = *C*_99_(*x*)) and *N* = 1000.

The eigenvalues plot in Fig. A1B also have complex conjugates; however, these points contain no new information as 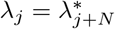.

### Appendix 4. Scaling of mode structure

Here, we show how both the localized modes and extended modes scale with discretization and friction. We claim that the extended modes are independent of the discretization scale; this is true unless N is small. Fig. A2 A shows how the number of extended modes on the basilar membrane scales with N. One can note that the number of these modes plateaus at *N* = 101. For values of *N* larger than this, 12 modes will remain for simulations run with our parameter values. This makes sense because for values of *N <* 101 the discretization scale *δx* is larger than the geometric wave number of the cochlea 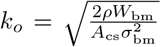 and the finite element model is a poor approximation. We can also see that increasing the discretization scale decreases the spacing between localized modes in Fig. A2 B,C. Note that some localized modes appear unstable in Fig. A2 B,C, this is a small N effect, Fig. A1 B shows the same system with *N* = 1000 where they are all stable.

We can also see how the discretization changes the behaviour for larger values of N. Here, we demonstrate that for low friction and large *N*, the localized modes approach the modes of a spatially uncoupled cochlea (grey dots showing eigenmodes of the matrix 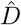), justifying the claim that in the low friction and large N limit localized modes act effectively independently. Fig A3 shows that if friction is 99% cancelled, increasing *N* makes localized modes qualitatively indistinguishable from the uncoupled modes. Note that they can not correspond exactly as there are 12 less localized modes than extended ones. We see a similar trend in Fig. A4 as we increase *f*. It is worth noting that even with only 1% of friction at resonance cancelled, the localized modes are not too dissimilar from the uncoupled modes as the fully passive cochlea already has low friction. If we make N larger at low *f*, we see a similar convergence, though localized and uncoupled modes are always further apart than an identical system with larger *f*.

### Appendix 5. Response Kernels

In the main text, we compared three different response kernels. Here, we expand on that comparison and show an additional response kernel and how changing the weighting of the zero derivative kernel affects the phase behaviour. The additional response kernel we investigate is a convolution of two exponentials,

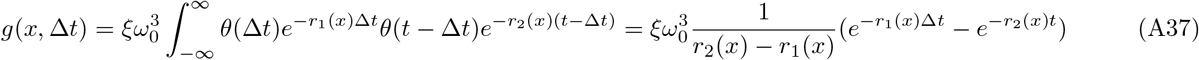

**Figure A2.**
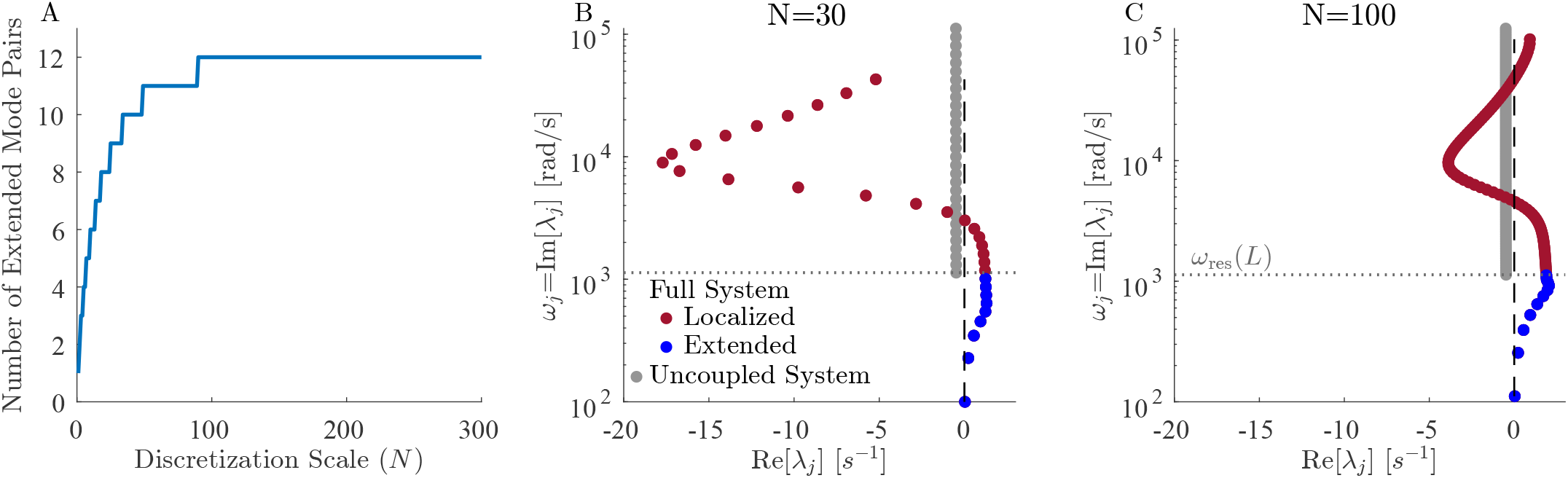
Modified Fig. 2 to show scaling with *N*. A) The number of extended modes as a function of discretization scale N. Note that this number plateaus at 12 starting at *N* = 101. B) Eigenvalue structure for N=30. C Eigenvalue structure for N=100. Note that the spacing between localized modes decreases as N is increased. B and C are identical to Fig. 2A except for the choice of N. These plots use exponential feedback with 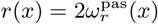 with friction 99% cancelled *C*(*x*) = *C*_99_(*x*).

**Figure A3.**
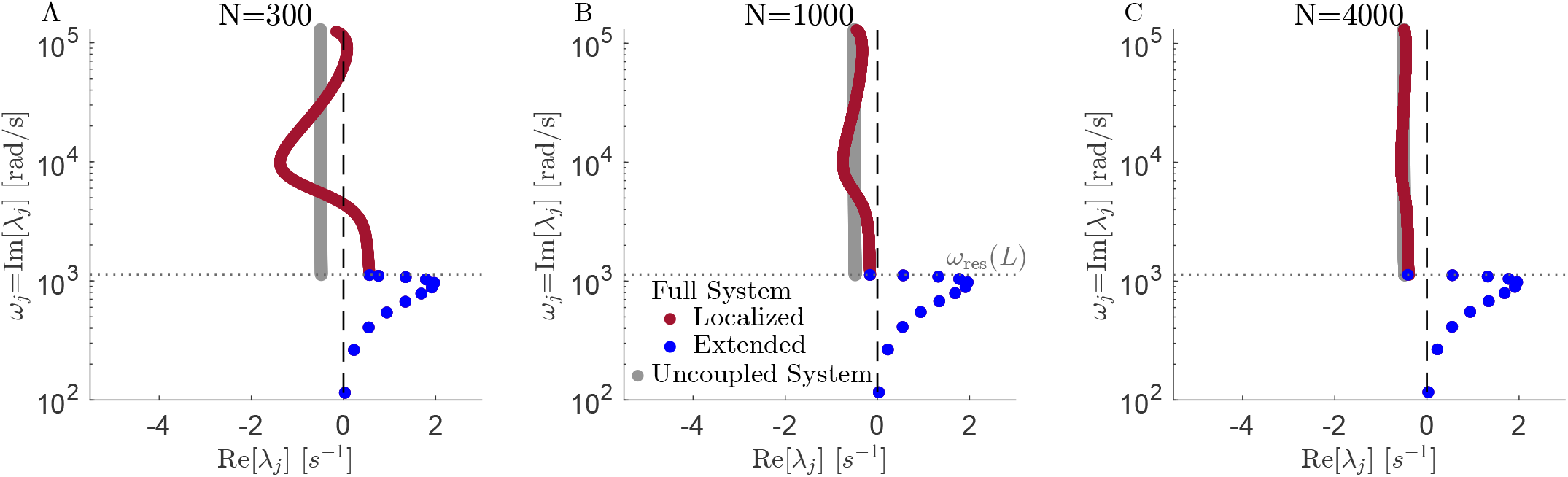
Modified Fig. 2A to show scaling with N. Comparison of the eigenvalue structure of localized modes with an uncoupled cochlea. Note that the localized modes become much closer to the uncoupled modes as we increase the discretization scale. A N=300. B N=1000. C N=4000. As N is increased, the localized modes approach the uncoupled system. Note that a discretization scale of 4000 has a spacing greater than that of the persistence length of the basilar membrane. These plots use exponential feedback with 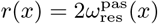 with friction 99% canceled (*C*(*x*) = *C*_99_(*x*)).

**Figure A4.**
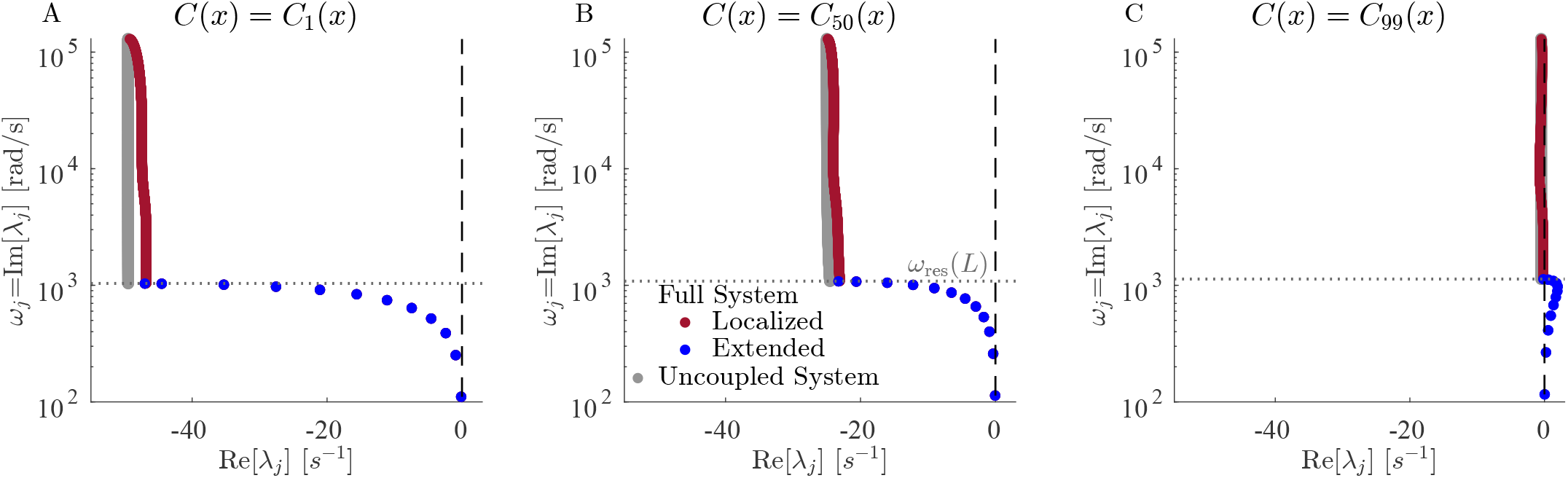
Modified Fig 2A to show scaling with friction. In this figure, we change *f* to see how this affects the distance of the localized modes from the uncoupled system. Note that the localized modes become closer to the uncoupled modes as we decrease the friction at resonance. a) 1% canceled. b) 50% canceled. c) 99% canceled. The cochlea is underdamped, so even with friction 1% cancelled, the localized modes are similar to the uncoupled ones. These plots use exponential feedback with 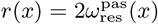 with friction f% canceled (*C*(*x*) = *C*_*f*_ (*x*)) and *N* = 1000.

**Figure A5.**
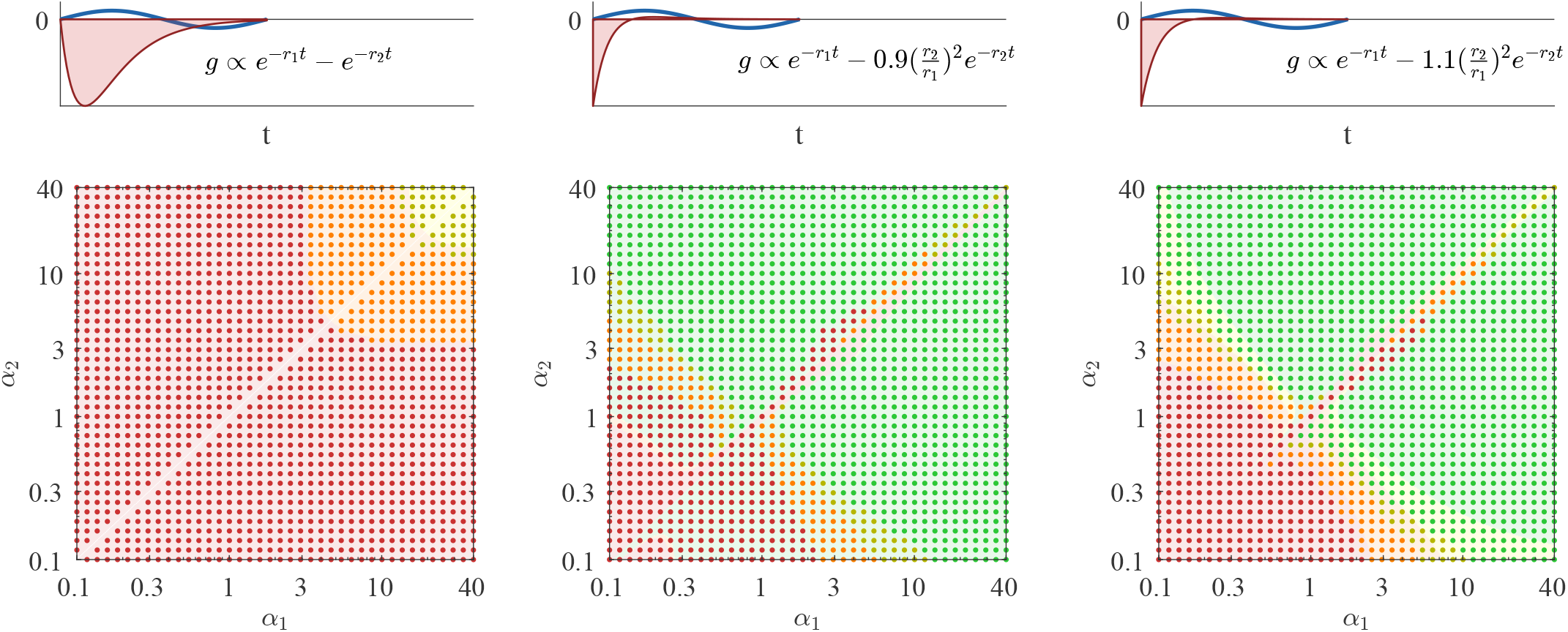
Changes to the response kernel *g*(*x*, Δ*t*) from Fig. 3. The middle and right columns add two exponentials, with two different choices of additive constant than that in Fig. 3 right. The left column shows two subtracted exponentials, equivalent to a convolution of two exponentials. The stability phase diagrams are analogous to Fig. 3 B. It is worth noting that the left column is qualitatively similar to a single exponential in that there is no stable region as *f* → 1 and that the left and right columns are no longer symmetric but do not deviate substantially from Fig. 3 B right.

One can show that this convolution simplifies to a sum with a relative prefactor of −1. The left column of Fig. A5 shows that this behaves in a way similar to an approximate derivative but without a region of true stability.

The other changes tried were to the response kernel,

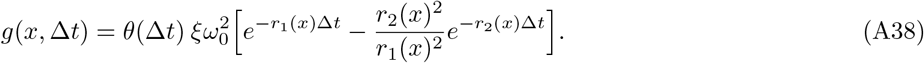

This kernel is motivated by having the property 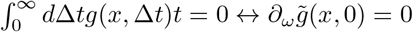. We found that it is stable for a large fraction of parameter space. To ensure this is not the product of fine-tuning, we change it so the derivative no longer integrates to exactly 0 in Fig A5. The phase diagram looks very similar except for some small changes close to the diagonal. The most important feature, stability for a large range of *α*_*j*_, remains true, indicating that our results do not require fine-tuning.

The other change to response kernels we tried was having a shift in the form of the response kernel,

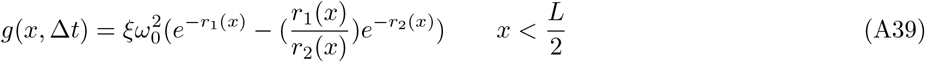

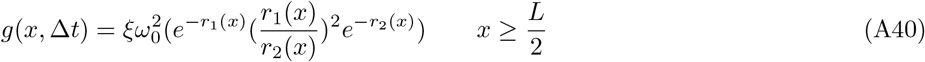

This phase behaviour is shown in Fig. A6, and we can see from the results that what is most important is the behaviour for the later half of the BM. This is expected since the latter half of the BM has lower resonant frequencies and would have a larger contribution to the extended modes.

### Appendix 6. Increasing Friction of the Oval Window

Two things can impact the stability of the extended modes. The first explained in the main text is the form of response kernel; more specifically, if the response kernel causes Im(*Z*(*x*., *ω*)) *<* 0 for values of *ω < ω*_res_(*x*). The imaginary part of impedance determines if an extended mode gains energy as it travels along the cochlea. The other not discussed in mian text is *ξ*_ow_; this is the amount of energy lost at the left end of the cochlea. Fig. A7 shows the behaviour of the three response kernels discussed in the main text when *ξ*_ow_ is multiplied by 100. One can see that eigenvalues are stable for a larger set *α*_*i*_ than before. This effect is particularly noticeable for larger values of *α*_*i*_. It is important to note that Fig A7 C,D are unchanged as the behaviour along the cochlea is unaffected by *ξ*_ow_. It is also worth noting that in Fig A7 E, we can see a large dip in the real part of localized modes that correspond to a frequency close to that of *ω*_ow_ this is once again an effect of increasing *ξ*_ow_.

**Figure A6.**
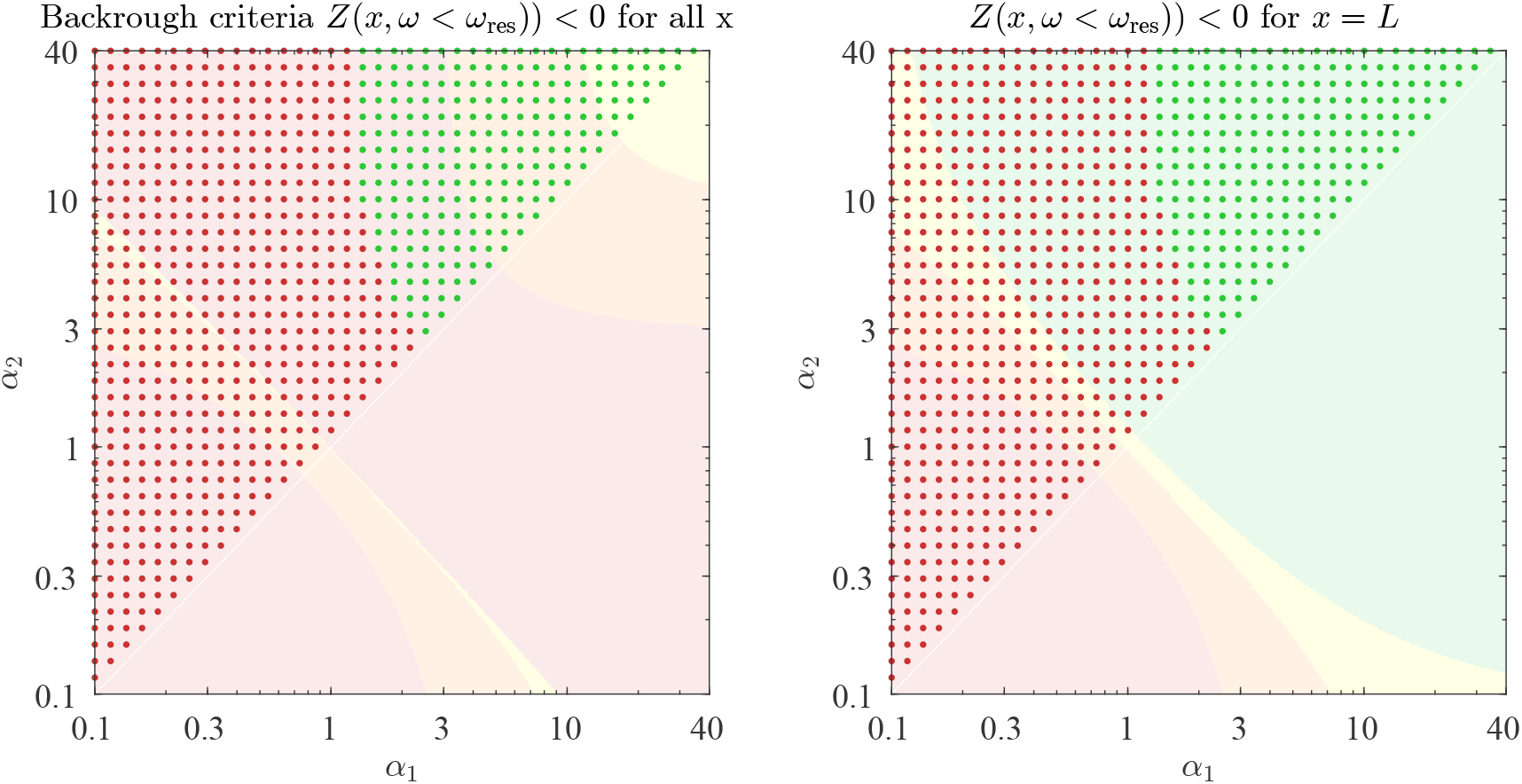
Combination of two response kernels from Eq. A39. Phase diagrams are plotted in the same style as Fig. 3B. In the left column, we use analytic criteria for instability that Ξ_net_(*x, ω*) *<* 0 for *ω < ω*_res_(*x*) for all values of *x*. While for the right column we use Ξ_net_(*x, ω*) *<* 0 for *ω < ω*_res_ only *x* = *L*. Neither criterion fully predicts simulation, unlike Fig. 3B; however, examining friction at *x* = *L* yields better predictions.

**Figure A7.**
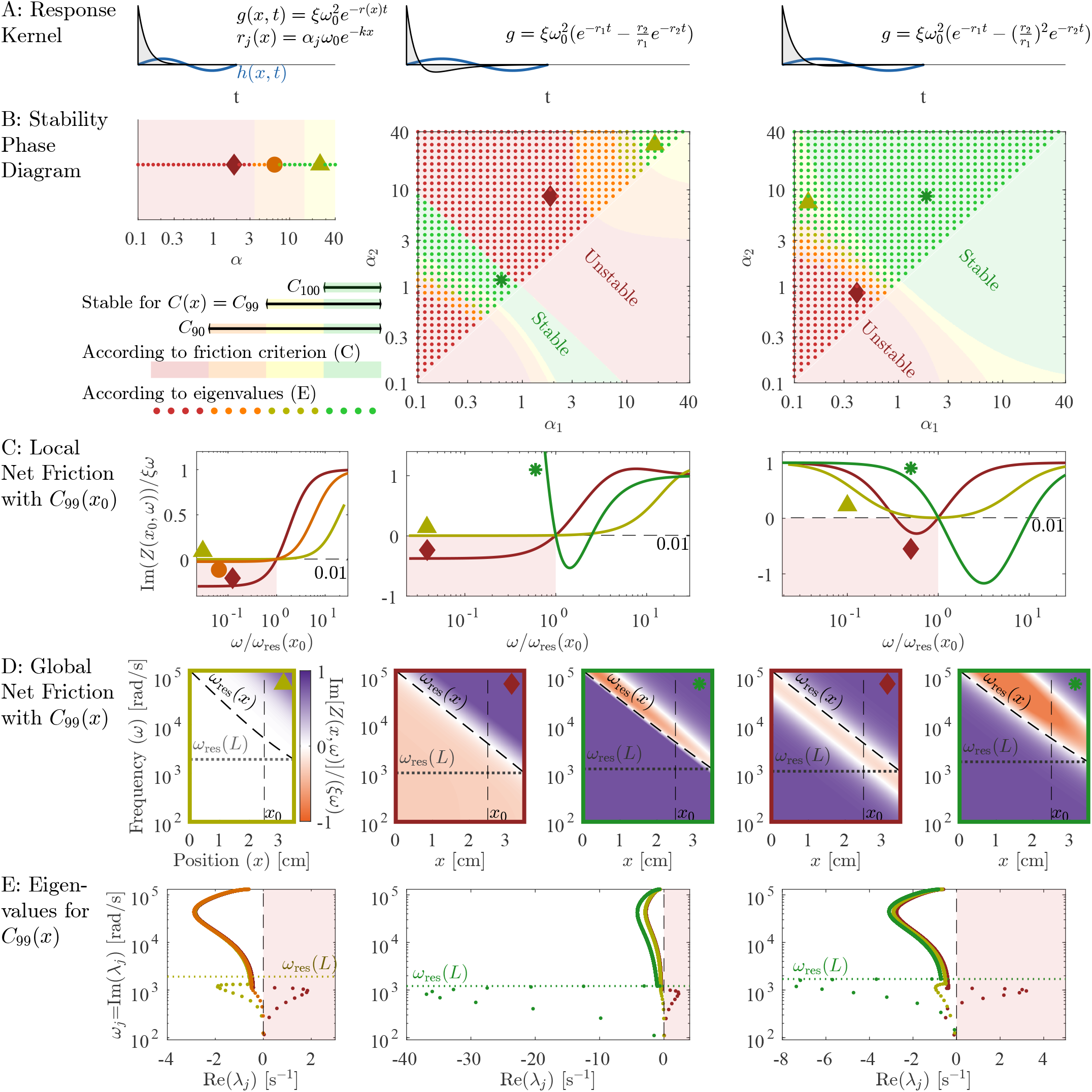
This figure is identical to Fig 3 in the main text except for now *ξ*_ow_ = 50000s^−1^, a value 100 times greater than normal. Points marked by large symbols are chosen to be the same as those from the main text, even if their stability should indicate a different colour. The criteria on Ξ_net_(*x, w*) is completely unchanged (background shading in B), only the eigenvalues are different. Note that at large values for *α*, the simulations shows stable points even if friction at low values of *ω* is negative.

